# DNA barcoding for parallelised single-molecule characterisation of kinetic phenotypes

**DOI:** 10.1101/2025.08.28.672975

**Authors:** Stefan H. Mueller, David M. Grimson, Holden Paz, Aidan C. Fitch, Haibo Yu, Marco Ribezzi-Crivellari, Andrew D. Griffiths, Antoine M. van Oijen, Lisanne M. Spenkelink

## Abstract

Single-molecule techniques provide exceptional resolution of biomolecular dynamics but are limited by low-throughput. We present a barcoding strategy that enables simultaneous kinetic profiling of multiple protein variants at the single-molecule level. Each protein is covalently linked to a unique DNA barcode, decoded via transient hybridisation of fluorescent probes distinguished by colour and binding duration. We applied this method to a library of 16 ClpS variants to systematically explore evolutionary trajectories towards a variant adapted to recognise N-terminal amino acids with binding kinetics optimised for single-molecule peptide sequencing. The approach uncovered ∼5-fold variation in median dissociation rates across variants. Several variants displayed bimodal kinetics, likely reflecting structural subpopulations, and differences in kinetic properties arose primarily from the relative abundance of these phenotypes. This approach offers a general framework for screening and optimising proteins for single-molecule applications, while revealing mechanistic insights that are inaccessible to traditional ensemble methods.

## Main

Single-molecule techniques have been used to probe ligand binding kinetics of protein binders and reaction kinetics in enzymes, revealing heterogeneous behaviours that are obscured in bulk measurements and transform our ability to probe the dynamics of biomolecular interactions^1-6^. These dynamics not only underpin molecular mechanisms but also play a critical role in biotechnological applications that use a dynamic read-out, such as single-molecule DNA^7-9^ and single-molecule protein sequencing^10, 11^. One way to enhance the performance of such technologies is to identify protein variants with more favourable kinetic properties. Achieving this requires the ability to assess large numbers of protein variants at the single-molecule level. However, most single-molecule methods can only screen a few variants at a time, limiting their capacity to explore large libraries or map sequence–function relationships at scale.

A major barrier to screening large protein libraries at the single-molecule level is the difficulty of identifying individual variants within a pooled assay. In typical single-molecule imaging, the number of distinguishable species is constrained by the limited number of fluorophores that can be used simultaneously. Some recent approaches, such as MUSCLE^12^ and SPARXS^13^, have attempted to increase throughput using indexing strategies, but these have so far been limited to nucleic acids or DNA-binding proteins. Tools for high-throughput single-molecule phenotyping of general protein libraries, however, remain underdeveloped.

Here, we describe a method that enables parallelised measurement of single-molecule phenotypes in protein libraries (Fig. 1a). Each protein is covalently linked to a DNA barcode via SNAP-tag chemistry^14^. These barcodes are read out through the transient binding of fluorescently labelled oligonucleotide probes. Combining the colour of the probe with the dwell time of its binding, allows the identification of hundreds to thousands of protein variants in the same assay. In the same experiment, we measure functional activity of these proteins.

**Figure 1.**
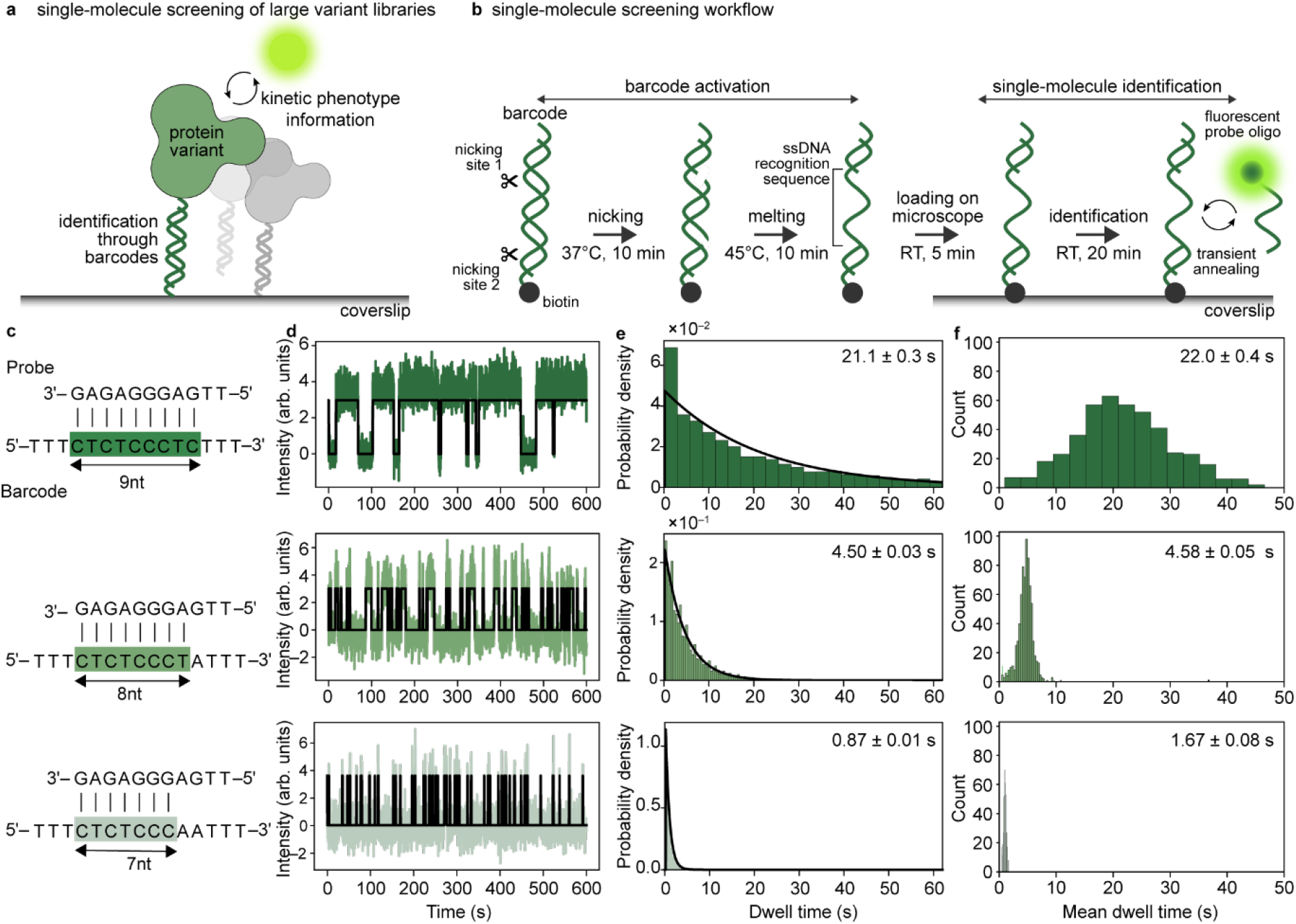
Single-molecule identification of variants. **a**, schematic representation of the method. Protein variants are attached to DNA barcodes, allowing unique identification at the single-molecule level. **b**, schematic representation of single-molecule identification using DNA barcodes with a single index. The recognition sequence (the index) is flanked by two nicking sites, allowing conversion of the recognition sequence to single-stranded DNA. After immobilisation on a microscope coverslip, fluorescently labelled probes transiently anneal to the barcode. **c**, barcode sequence design showing a single probe with complementarity to three index sequences. **d**, example single-molecule trajectories of probe binding, showing intensity as a function of time, with Hidden-Markov-Model fit shown in black, for the 9-nt (top, N=5600 dwell times), 8-nt (middle, N=22693 dwell times), and 7-nt (bottom, N=13181 dwell times) complementary sequences shown in a. **e**, dwell-time distributions corresponding to each barcode sequence. The black line represents a single-exponential fit to the data, giving characteristic time scales of 21.1 ± 0.3s, 4.50 ± 0.03s, and 0.87 ± 0.01s for the 9-nt (top), 8-nt (middle), and 7-nt (bottom) complimentary sequences, respectively. **f**. Histograms of mean dwell times per molecule for the 9-nt (22.0 ± 0.4 s, N=441 barcodes), 8-nt (4.58 ± 0.05 s, N=750 barcodes) and 7-nt (1.67 ± 0.08 s, N=467 barcodes) complimentary sequences, respectively. Errors represent S.E.M.

To showcase the power of this approach, we apply it to a library of 16 variants of Planctomycetes ClpS^15-17^, a member of the ClpS N-end rule adapter family that is used to identify N-terminal amino acids in single-molecule protein sequencing^10^. Optimal sequencing depends on binding kinetics, rather than affinity, since repetitive probing via rapid on-and-off binding is central to accurate amino acid identification. Our assay quantifies real-time binding kinetics of each variant and identifies a single mutation that alters peptide association rates by a factor of five. Several variants displayed bimodal kinetics, with differences in kinetic behaviour driven primarily by the relative abundance of each kinetic phenotype. These findings highlight the ability of kinetic measurements to detect phenotypes that traditional equilibrium-based methods would miss. This scalable platform should enable the power of single-molecule techniques to be applied for high-throughput phenotyping of protein libraries.

## Results

### Single-molecule sequencing by hybridisation

We aimed to develop an identification method, based on labelling protein molecules with DNA barcodes, that can be scaled to measure single-molecule ligand-binding kinetics of hundreds or thousands of different protein molecules within the same experiment (Fig. 1a).

To demonstrate this concept, we focused first on very simple barcodes with a single recognition sequence (index) (Fig 1b). We designed a fluorescently labelled probe and corresponding barcodes with index sequences with 9-nt, 8-nt, and 7nt complementarity to the probe (Fig. 1c). The recognition sequence is flanked by two nicking sites. After nicking, the short, nicked fragment is melted off by incubation at 45°C, to expose the single-stranded recognition sequence. This activated barcode, which has a biotin at the 5′ end, is then immobilised on a streptavidin-coated microscope coverslip in a microfluidic flow cell. We then introduce the fluorescently labelled probe oligonucleotides and monitor probe hybridisation to the indices. Since the region of complementarity between the probe and the indices is very short, annealing of the probe to the barcode is transient. Furthermore, with these short sequence lengths, probe binding depends on the number of complementary nucleotides between the index and probe, whereby a probe binds a 7-nt index >20 times shorter than a 9-nt index^18^. Therefore, the dwell time can be used to identify the index length and, in turn, the barcode identity. By measuring fluorescence intensity as a function of time for each barcode, we generate single-molecule trajectories (Fig. 1d). From these trajectories, we can extract the duration of each binding event (the dwell-time). As expected, these events follow an exponential dwell-time distribution (Fig. 1e), while the distribution of mean lifetimes across molecules follows a normal distribution, consistent with the central limit theorem (Fig. 1f). As expected, single-molecule trajectories (Fig. 1d) and the resulting dwell-time distributions (Fig. 1e,f) decreased with the length of the index complementary to the probe, passing from 21.1 ±.0.3 s for 9-nt, to 4.50 ±0.03 s for 8-nt and 0.87 ± 0.01 s for 7-nt. This large difference in dwell time between the three sequences, allows us to identify the index.

Because the variance of an exponential distribution grows with the square of the mean, the resulting normal distributions also broaden for longer-lived interactions (Fig. 1f). This inherent scaling sets a practical upper limit of approximately 20 seconds for mean lifetimes—beyond which accurate classification would require prohibitively long imaging times.

### Scaling barcode diversity through multi-index design

To increase the number of identifiable variants, we extended our approach to barcodes comprising two indices (Fig. 2a), where each index is recognised by a different spectrally distinct probe. Each index sequence is chosen from a set of three, with distinct probe-hybridisation lifetimes, allowing independent identification of both indices from the same field of view. By combining two orthogonal indices—each selected from a set of three, with distinct probe-hybridisation kinetics—we generate 3×3=9 unique barcode combinations. During imaging, the hybridisation kinetics of each index are measured independently in their respective fluorescence channels, before colocalisation of the signals (Fig. 2b). Each barcode, therefore, maps to a distinct pair of mean dwell times, forming a cluster in two-dimensional space. Barcode identity is then determined in an unbiased and fully automated fashion using a clustering algorithm (Fig. 2c; see Methods). To evaluate the accuracy of our classification method, we repeated the experiment using a subset of six two-index barcodes. We then counted how often the algorithm identified a barcode that was not actually present. We achieved 95% accuracy (Extended Data Fig. 1) and obtained over 170 correctly identified molecules for each barcode. The remaining 5% misclassification is unlikely to significantly impact downstream analyses.

**Figure 2.**
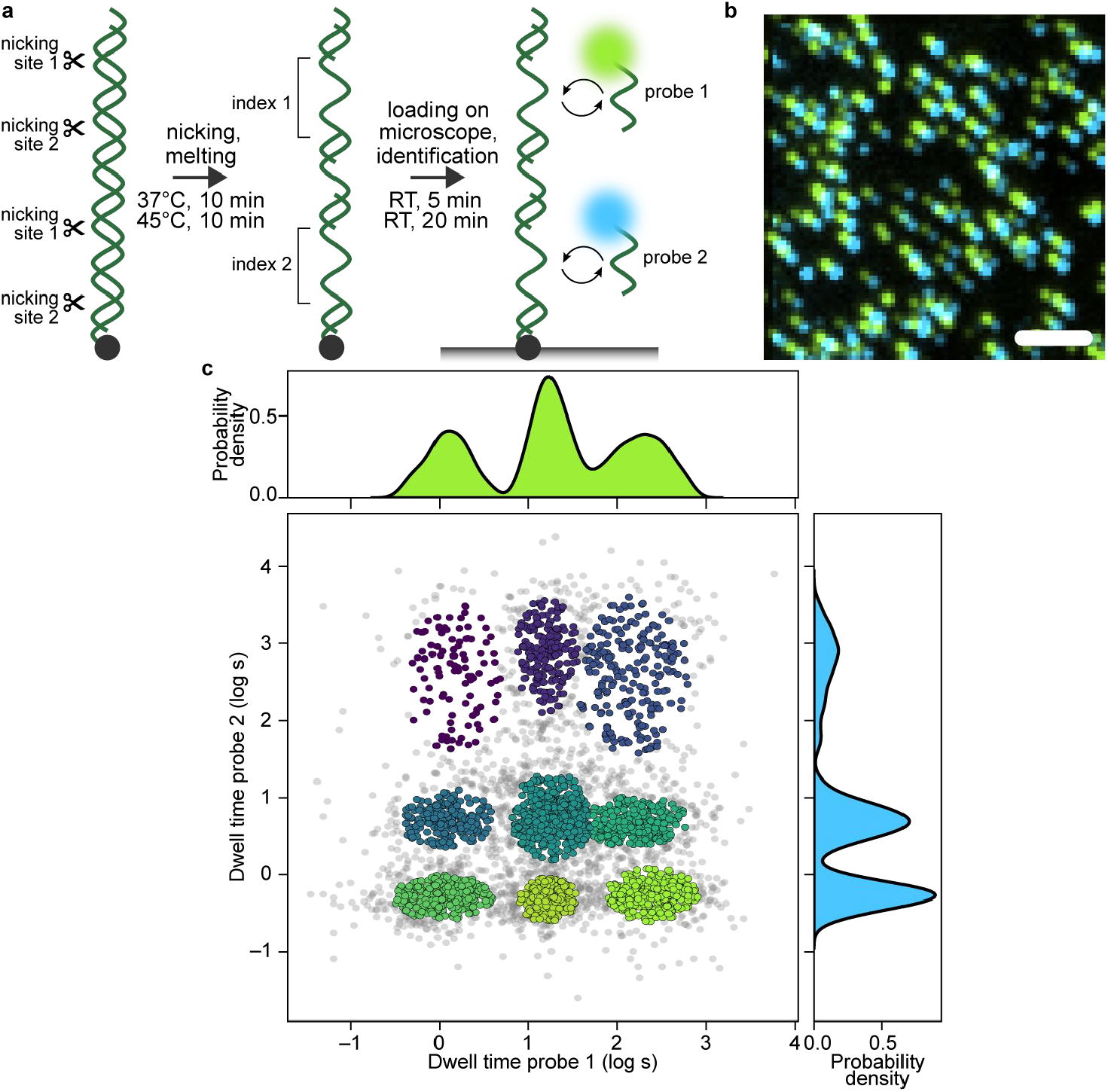
Increasing the number of barcodes. **a**, schematic representation of the 2-index barcodes. Barcode activation and detection is carried out as described for 1-index barcodes. Each index binds a spectrally different fluorescent probe. **b**, standard-deviation projection of a representative field of view showing colocalisation of signals for probe 1 (light green) and probe 2 (dark green, offset by 4 pixels). **c**, hybridisation kinetics for nine two-index barcodes. Combined measurement of mean dwell times for probe 1 (top) and probe 2 (right) generates nine distinct clusters (middle), where each dot represents a single two-index barcode. The grey spots are more than 1.5 standard deviations from the cluster mean and discarded to increase the fidelity of barcode identification.

To increase the combinatorial capacity of our barcoding strategy, we designed and tested six orthogonal probe sequences, each capable of binding three indices with distinct hybridisation kinetics (Extended Data Fig. 2, Extended Data Tables 1 and 2). The probes are labelled in three colours, with two probes per colour. Because spectrally distinct probes can be loaded simultaneously, we require two rounds of imaging to measure all six probes. This yields a total of 6 ×3 = 18 unique indices.

### Simultaneous single-molecule profiling of ClpS variants

To demonstrate the utility of our barcoding approach for parallel functional profiling, we applied it to a 16-member library of ClpS variants. Planctomycetes ClpS is an N-end rule adaptor protein^15, 16^ used in a single-molecule protein sequencing platform^10^, where kinetic properties (the association and dissociation rate constants, *k*_on_ and *k*_off_), rather than thermodynamic properties (the equilibrium constant, *K*_D_), determine sequencing performance. Fast kinetics enable rapid turnover and time-resolved signal detection, which are critical for sequencing accuracy.

A ClpS variant previously developed for single-molecule protein sequencing^10^ (ClpS-Opt) contains four mutations relative to the wild-type ClpS (WT)^17^: Q55R, L68M, V72M, and Y100R^10^. This variant was optimised for recognition of N-terminal leucines in peptides. To systematically investigate the functional contribution of each mutation, we constructed a complete combinatorial gene library (Fig. 3a) comprising all 2^4^ = 16 possible variants across these four positions, including WT and ClpS-Opt (Fig. 3b).

**Figure 3.**
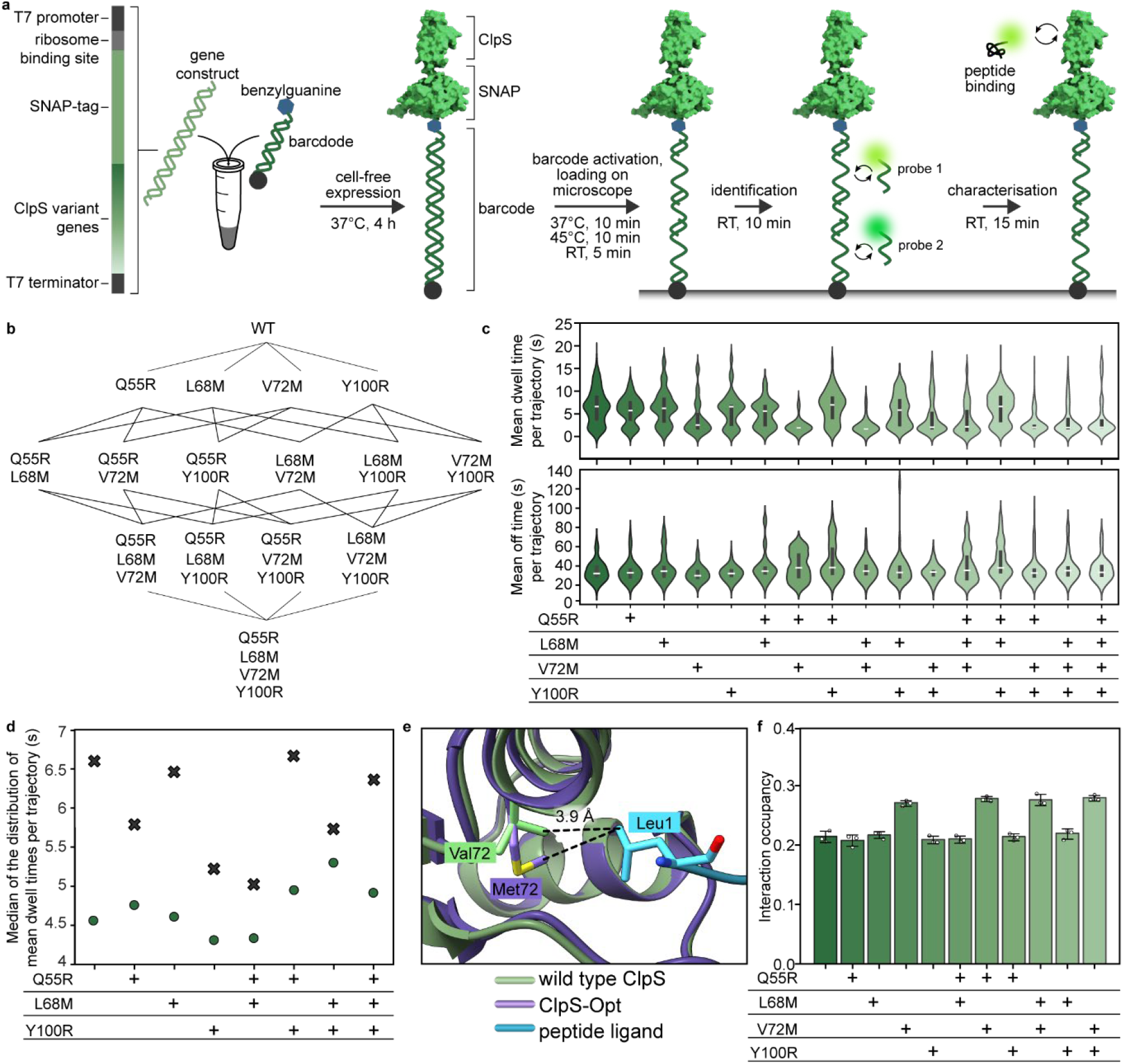
Simultaneous measurement of peptide binding kinetics for 16 ClpS variants. **a**, Schematic representation of the workflow. Gene constructs encoding for ClpS variants fused to a SNAP-tag are expressed in a cell-free expression system. After expression, the SNAP-tag covalently binds to the barcode. After barcode activation, all barcoded constructs are loaded onto the coverslip, followed by identification and characterisation. **b**, sequence space between wild-type ClpS and ClpS-Opt. **c**, violin plots showing the distributions of the mean dwell times per trajectory (top) and mean off time per trajectory (middle) for all 16 ClpS variants. The black bars correspond to the interquartile range (25th to 75th percentile), and the white lines mark the median. **d**, medians of the dwell times distributions of the mean dwell time per trajectory in c, comparing variants with (green circles) and without (black crosses) the V72M mutation. **e**, AlphaFold2 structures of the wild-type (green) and V72M variant (purple) of ClpS, bound to the peptide (cyan). **f**, Occupancies of the van der Waals contacts between the terminal leucine residue with ClpS Val/Met72, determined using the ProLIF molecular-dynamics package. Each data point represents a single replicate showing the percentage of the 50,000 trajectory frames that have a van der Waals contact between the two residues of interest. The error bars represent the standard deviations.

To characterise phenotypes across the ClpS library simultaneously, we covalently linked each ClpS variant to a unique DNA barcode using SNAP-tag chemistry^14^. Gene constructs encoding SNAP–ClpS fusion proteins were expressed in a cell-free system^19^, in the presence of a barcode. The expressed SNAP-tag (O^6^-alkylguanine DNA alkyltransferase) reacts irreversibly with a benzylguanine on the 3′-end of the DNA barcode, to establish a covalent link between the protein and the DNA barcode (Fig. 3a). As before, DNA barcodes are biotinylated at the 5′ end to allow immobilisation of the barcoded SNAP–ClpS proteins on a microscope coverslip.

Following simultaneous immobilisation of all 16 variants, two spectrally distinct probe oligos were introduced to enable barcode identification (Fig. 3a). After identification, a fluorescently labelled peptide containing an N-terminal leucine was added to monitor binding to the ClpS variants. We quantified peptide association and dissociation kinetics across all 16 variants in parallel (Fig. 3c). Fig. 3c shows the distribution of the mean dwell time (top) and mean off time per trajectory. Along the possible evolutionary trajectories from wild-type ClpS to ClpS-Opt, we observed a ∼5-fold reduction in the median of these dwell time distributions.

Furthermore, we observed bimodal distributions in the dwell times of several variants, including wild-type ClpS. To confirm that this was not due to barcode misclassification or colocalisation of distinct barcodes, we analysed wild-type ClpS by itself (Extended Data Fig. 3). This measurement recapitulated the bimodal distribution. The fast and slow modes of the mean dwell time distributions occur at similar values across all variants (i.e., where the violins are widest). (Extended Data Fig. 4a,b). These observations suggest the presence of two subpopulations of the ClpS–peptide complex, with the mutation state influencing their relative abundance. Indeed, the differences in the median of the mean dwell times are principally due to differences in the relative abundance of these two subpopulations (Extended Data Fig. 4c), wild-type ClpS being dominated by the slow-off subpopulation and ClpS-Opt by the fast-on, fast-off subpopulation.

The V72M mutation had the largest effect on peptide binding kinetics. Figure 3d shows a comparison of the mean dwell times variants with or without the V72M mutation. Remarkably, some double and triple mutants comprising V72M exhibit less dramatic effects than V72M alone, even though the same mutations without V72M decrease dwell times, suggesting complex epistatic relationships.

To better understand the molecular basis of these kinetic changes, we performed molecular dynamics simulations of ClpS–peptide interactions (Fig 3e,f). The simulations revealed altered van der Waals (Fig. 3e) and electrostatic interactions (Extended Data Fig. 5a,b) in variants containing the V72M mutation with the N-terminal leucine residue of the peptide. Consistent with our experimental observations, the other mutations did not induce significant changes in these interactions.

## Discussion

We have developed a DNA barcoding workflow for high-throughput, single-molecule identification and functional profiling of protein variants. We applied this approach to quantify peptide binding kinetics of 16 variants of the Planctomycetes ClpS protein in a single experiment. These variants cover all possible evolutionary trajectories between wild-type ClpS and an engineered variant, ClpS-Opt, optimised for single-molecule protein sequencing^10^.

While all mutations and combinations thereof had some effect on binding kinetics, and there was evidence of complex epistatic interactions, we find that the V72M mutation dominates changes in peptide binding kinetics. Methionine is typically considered chemically inert and structurally passive^20^, making this observation unexpected. However, consistent with our experimental results, our molecular dynamics simulations revealed increased van der Waals and electrostatic interactions between residue 72 and the peptide’s N-terminal leucine.

Furthermore, our single-molecule read out revealed two kinetic states: a slow-off state with a dwell time of ∼7 s, and a fast-off state with a dwell time of ∼1.5 s, with the overall median dwell time largely governed by the relative frequency of these two states. This highlights a central strength of single-molecule techniques: the ability to uncover functional changes in molecular dynamics that bulk thermodynamic assays or simulations may overlook^1, 2, 21, 22^. In wild-type ClpS and other variants that do not contain the V72M mutation, both states are present, whereas in variants with the V72M mutation the fast state predominates. In the cell, ClpS binds N-terminal residues with affinities that are modulated by its interaction with ClpA^23^. This dynamic regulation may involve changes in dwell time during substrate engagement and transfer, conceptually similar to the switching we observe between the two kinetic states.

A key advantage of simultaneous measurement is the elimination of experiment-to-experiment variation, which is critical in single-molecule assays, where experimental conditions can be difficult to standardise^24, 25^. In addition, parallelisation reduces measurement time substantially. In our case, measuring 16 variants in parallel yields an effective 16-fold reduction in acquisition time per replicate, and a total 45-fold (16 × 3 – 3) reduction for triplicates. Beyond measurement efficiency, our workflow also offers practical advantages. Gene variants were ordered as synthetic eBlocks, eliminating the need for cloning and simplifying construct preparation. Proteins were expressed via cell-free expression, and barcode coupling occurred in the same reaction through SNAP-tag chemistry, avoiding extra purification steps. These features streamline assay setup and make the method well-suited to rapid, iterative experiments.

While we demonstrated our method using 16 variants, it is inherently scalable. To project the potential number of uniquely identifiable barcodes, we calculated the total number of index combinations, using the binomial coefficient with replacement. This yields 171 combinations for barcodes with 2 indices. If we add just one more index per barcode, this number increases to 969. When we use six indices per barcode, we can uniquely identify more than 100,000 variants. This capacity enables screening at the scale of directed-evolution campaigns (10^5^–10^8^ variants), proteome-wide studies (20,000–40,000 human proteins)^26-28 29^, and deep mutational scanning of individual proteins.

Future improvements could further expand this capacity. Our current implementation uses fixed probe sets and discrete kinetic bins. Applying machine learning to binding trajectories could allow detection of more subtle kinetic differences. Introducing mismatches into index sequences would further increase barcode diversity without expanding the probe set.

Another method developed over 10 years ago, SMI-seq, used halo-tagging to attach barcodes^30^. However, identification in this method relied on *in-situ* amplification and sequencing. Other high-throughput single-molecule methods, such as SPARXS^31^, MUSCLE^12^, and magnetic-bead-based hairpin unzipping^32^, have been applied to screen nucleic acid variants. However, these platforms are limited to measuring DNA variants (*e.g*., different sequences or secondary structures) and the effect of enzymes, such as Cas9, on these DNA variants. By contrast, our SNAP-tag approach enables direct measurement of protein phenotype. SNAP tags have been successfully used to tag a plethora of proteins ^33-36^, suggesting that our DNA barcodes are compatible with these proteins. Our system could also be integrated with SNAP display workflows, in which large libraries of gene variants are expressed *in vitro* in emulsion droplets and covalently linked to the proteins they encode via SNAP-tag chemistry^37, 38^. Encoding barcodes directly into gene constructs would allow single-molecule screening of large libraries in directed evolution experiments.

We have also developed an alternative strategy for parallelised single-molecule analysis based on peptide barcoding. This approach uses genetically encoded peptide tags that are decoded directly using single-molecule protein sequencing after single-molecule analysis of ligand-binding kinetics^39^. While this alternative approach avoids the need to couple DNA barcodes to proteins, it requires proprietary hardware and chemistry. Our DNA barcodes are compatible with standard TIRF microscopy and off-the-shelf reagents, making it broadly accessible and adaptable. Furthermore, use of SNAP-tag barcoding and IVTT expression fits neatly into emulsion-based workflows for library screening.

In summary, our barcoding method provides a generalisable, scalable strategy for single-molecule kinetic profiling of proteins. It bridges the gap between high-throughput sequencing-based platforms and protein functional analysis, opening new directions for proteomics, protein engineering, and directed evolution.

## Methods

### Preparation of DNA-barcodes

Single-stranded 150 nt DNA oligomers were synthesised by IDT (Ultramers, IDT). To introduce the end-modifications, the oligomers were amplified using the polymerase chain reaction (PCR) reaction (Q5 Hot Start NEB) with biotin/benzylguanine-modified primers (Barcode_FW, Barcode_RV, Extended Data Table 3) for 30 cycles under the following conditions: 10s at 98°C, 20s at 64°C, 60s at 72 °C followed by a final extension for 120s at 72 °C. Finally, the amplified DNA was purified using silica-membrane spin-columns (Wizard SV, Promega).

### Preparation of gene-constructs

For all ClpS constructs, the full SNAP-fusion construct was ordered as a gene fragment (eblocks, IDT, for full sequences see Extended Data Table 4) and amplified by PCR reactions (Q5 Hot Start NEB) under standard conditions as described by the manufacturer with the primers RecA_FW and RecA_RV (Extended Data Table 3). PCR products were purified using Wizard® SV Gel and PCR Clean-Up System (Promega) using conditions described by the manufacturer. Purified products were stored at – 20°C.

### Expression of barcoded variants

All proteins were expressed by *in vitro* transcription and translation (IVTT) using the PURExpress *in vitro* protein synthesis kit (NEB) as per the manufacturer’s instructions. Only 15 µL reactions were required for ample sample for single-molecule characterisation. In addition to the variant gene and the kit components, reactions also contained 60 ng of DNA barcodes (with biotin and benzylguanine modifications), murine RNAse inhibitors (NEB) and disulfide bond enhancers (NEB) as per manufacturer’s instructions. Reactions were incubated at 37°C for 4h for protein expression and subsequently 12-15h at 4°C to allow coupling of the DNA barcode to the SNAP-tagged protein. Note that all components were stored in DTT-free and EDTA-free buffers to avoid inhibition of the SNAP-reaction. After incubation 1 µL of IVTT was mixed with 1 unit of nicking endonucleases Nt.BsmAI and Nb.BtsI respectively, in 10 µL Cutsmart buffer (NEB). After vigorous mixing with a pipette, the reaction was incubated for 10 minutes at 37°C to allow for nicking, before 20-fold dilution in phosphate-buffer saline (PBS) buffer and incubation at 45°C for a further 10 minutes. This resulting sample could either be used for single-molecule microscopy immediately or stored at –80°C for later use.

### Preparation of fluorescently labelled peptide

A peptide (LAKLDEESILKQC) with an N-terminal leucine for recognition by ClpS and a c-terminal cysteine for labelling was purchases from Genscript. The peptide was dissolved at 1 mg/mL in labelling buffer (20 mM HEPES, pH 7.6, 1 mM EDTA) and reduced by addition of 5 mM tris(2-carboxyethyl)phosphine (TCEP) for 1 h at room temperature. Alexa Fluor 647 C2 maleimide (Invitrogen) was dissolved in anhydrous DMSO to a final concentration of 10 mM, and added to the reduced peptide in a 10-fold molar excess. The reaction was incubated for 2 hours at room temperature, protected from light. Unreacted dye was removed using a Zeba spin desalting column (Thermo Fisher) equilibrated in labelling buffer. The degree of labelling and peptide concentration were determined by absorbance at 280 nm and 650 nm using a NanoDrop spectrophotometer.

### Single-molecule microscopy

First, we prepared microfluidic flow chambers as described before^36, 40, 41^. Briefly, cover slips (24 × 24 mm, Marienfeld) were functionalised with biotin-PEG (Laysan Bio). A polydimethylsiloxane (PDMS) block, with 1 mm wide and 0.1 mm high channels, was made using soft-lithography^42^ and placed on top of the cover slips. Two stretches of polyethylene tubing (PE-60: 0.76 mm inlet diameter and 1.22 mm outer diameter, Walker Scientific) were inserted into the PDMS block at both ends of the channel and connected to a syringe pump (Adelab Scientific) to allow for buffer flow. The flow-cell device was then mounted on an inverted total-internal reflection fluorescence (TIRF) microscope (Nikon Eclipse Ti-E), with a temperature-controlled stage (25°C; Okolab), a 100x TIRF objective (NA = 1.49, oil, Nikon) and an EMCCD camera (Andor iXon).

Before the start of experiments, the flow channel was incubated with blocking buffer (50 mM Tris–HCl pH 7.6, 50 mM Potassium Chloride, 2% (v/v) Tween-20) to minimise nonspecific binding of DNA or proteins to the cover-slip surface. Barcoded protein-variants were then immobilised using a flow of 70 µL/min for 100µL followed by an additional flow of 5 µL/min for 80µL.

### Phenotyping reactions

To initiate the phenotyping reaction 20 nM of fluorescently labelled peptide was diluted in phenotyping-buffer (PBS supplemented with 10 mM ascorbic acid, 1 mM UV-aged trolox, 2.5mM protocatechuic acid (PCA) and 50 nM protocatechuate-3,4-dioxygenase (PCD), 0.0025% Tween-20, 0.04 mg/mL BSA). The sample was then loaded at 70 µL/min for 100 µL. A constant flow rate of 2 µL/min was maintained for the duration of the measurement (10min).

The sample was illuminated using a 532-nm laser (Coherent, Sapphire 532–200 CW) at 5.2 mW cm^−2^. If not otherwise specified, a frame rate of 5 frames per second with an exposure time of 200ms was used.

### Barcoding reactions

To initiate the barcoding reactions 10 nM of two different probe oligos labelled with the fluorescent dyes Alexa Fluor 488 or Alexa Fluor 647, respectively, were loaded in barcoding buffer (25 mM Tris-HCl pH 7.6, 250 mM Magnesium acetate, 0.0025% Tween-20, 0.04 mg/mL bovine serum albumin (Sigma), 0.2 mM EDTA, 10 mM ascorbic acid, 1 mM UV-aged Trolox, 2.5 mM protocatechuic acid, 50 nM protocatechuate-3,4-dioxygenase).

Samples were imaged for 10 minutes, consecutively using a 488 nm laser (Coherent, Sapphire 488–200 CW) at 1.6 mW cm^−2^ and a 647-nm laser (Coherent, Obis 647–100 CW) at 5.2 mW cm^−2^ respectively. If not otherwise specified, a frame rate of 5 frames per second with an exposure time of 200 ms was used.

### Data analysis

Data analysis was carried out using programs written in Fiji^43^ and Python. The raw data was first corrected for a non-uniform excitation-beam as previously described^25^. To correct for mechanical drift of the microscope stage during the measurement we tracked the position of a fluorescent bead within the field of view which is visible for the entire movie by the means of Gaussian peak fitting.

Next, to detect all fluorescent spots corresponding to probe-hybridisation, we calculated the average intensity value for each pixel and detected peaks using a threshold approach as described before ^25^. Finally, we integrated the intensity of the selected peaks over time and applied a local background correction ^25^.

To characterise hybridisation kinetics, we first assigned bound and unbound states in the trajectories by the use of Hidden-Markov models (HMM) as described by Baum and Petrie^44^. The fluorescence intensity of the bound and unbound states can be approximated by Gaussian distributions, we therefore utilize a HMM with Gaussian emission probabilities implemented in python^45^. We estimate the optimal parameters of the HMM via the Baum-Welch algorithm. To avoid convergence in local extrema we determine an initial guess by fitting the distribution of all intensity values at all times to a bi-normal distribution as previously described^46^. Finally, we compute the most likely state-transitions for all individual trajectories using the Viterbi-algorithm^47^.

### *In silico* modelling, molecular dynamics, and relative binding free energy simulations

All molecular dynamics simulations were carried out using Amber22^48^. Three independent replicates for each system were simulated, totaling 15 μs for each apo- and peptide-bound system. Wild-type ClpS, variants involving a single amino acid substitution, and variants involving two simultaneous amino acid substitutions were simulated. Systems were set up using CHARMM-GUI^49, 50^. The intrinsically disordered region (residue 1 to 28) was truncated from the model. The N-terminus and C-terminus were capped with a proline N-terminal group (PROP) and an N-methylamide (CT3) group, respectively. The ff14SB protein force field was used for proteins^51^. The TIP3P model was used for water^52^.

Molecular dynamics simulations were performed after solvating the system in a cubic box with a size 37 Å^50^, or at least 10 Å from the solute interface. Na^+^ and Cl^−^ counter ions were added to neutralise the system and achieve a salt concentration of 0.15 M. Simulations were performed using periodic boundary conditions (PBC) at constant temperature (303.15 K) with the Langevin thermostat (a damping coefficient of 1 ps^-1^)^53, 54^ and at a pressure of 1.0 bar using the Monte-Carlo (MC) barostat^55^. Short-range electrostatics were calculated together with long-range electrostatics particle mesh Ewald (PME) with a cut-off of 9.0 Å^56^. For all systems, energy minimisation (5000 steps with 2500 steepest-descent steps) and 125 ps equilibration with a 1 fs timestep were performed first with positional restraints placed on all the protein-heavy atoms (with a force constant of 1.0 kcal/mol/Å^2^ on the protein atoms. Initial velocities were taken from a Maxwell distribution at 303.15 K. The equilibration was followed by 5 μs production runs for the apo and peptide-bound proteins, respectively. Hydrogen mass repartitioning (HMR) was applied with the time step set to 4.0 fs^57^, and all covalent bonds involving hydrogens were kept rigid with the SHAKE algorithm^58^. Snapshots were saved every 100 ps. VMD (Visual Molecular Dynamics^59^), CPPTRAJ^60^, and UCSF ChimeraX^61^ were used for analysis and visualisation.

Alchemical free energy calculations based on thermodynamic integration enable the estimation of the relative binding free energy of two ClpS+peptide systems (for example, WT and V72M) through a thermodynamic cycle (Extended Data Fig. 6). Relative binding free energy differences for the model peptide binding to ClpS and the V72M variant were calculated using eq 1:

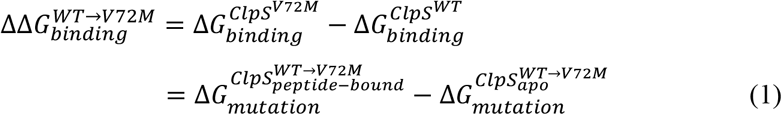

A detailed description of the method can be found elsewhere^62^.

Thermodynamic integration simulations were initialised with independent snapshots taken from molecular dynamics simulations. The starting structure was mutated at a single residue with MODELLER^63^, and single topologies were prepared using tleap and the timerge module of parmed^50^. Each alchemical transformation involved 21 evenly spaced values of λ (0, 0.05, … 0.95, 1). Bonds containing hydrogen atoms were constrained with the SHAKE algorithm except those in the alchemical region. The gti_add_sc value was set to 5. Energy minimisation was performed by evolving the system at 1 K for 125 ps at each λ window, and initial velocities were taken from a Maxwell distribution at 303.15 K. The minimised structure was then heated at 303.15 K for 125 ps in the NVT ensemble. All protein residues were restrained with a weight of 10.0 kcal/mol/Å^2^ during minimisation and 1.0 during the heating step. The timestep was set to 1 fs. The rest of the simulation parameters are identical to those used in the molecular dynamics simulations. NPT trajectories were then collected with 10 ns per λ window. For the softcore potential^64^, the second-order smooth step potential was used, and values of α and β were 0.2 and 50 Å2 as recommended^62^. There was no net charge in the alchemical transformation, and any correction for electrostatic artifacts was expected to be negligible. The first 25 ps of collected λ-derivative of the total energies were discarded as equilibration, and the remaining data were integrated to calculate the ΔG values for each alchemical transformation. Results from independent snapshots of the initial and final apo and peptide-bound states were averaged to mitigate initial structure-dependence.

Structural prediction was performed with the ColabFold implementation of AlphaFold2^65^. The source code was retrieved from the GitHub repository (https://github.com/YoshitakaMo/localcolabfold, access date: 05.23.2024). For each prediction, five models were generated, and the top model was used as the initial structure for molecular dynamics simulations.

## Data availability

The data generated in this study are available in the Zenodo repository under DOI: 10.5281/zenodo.16149314.

## Code availability

The custom scripts are available in a public GitHub repository at https://github.com/Single-molecule-Biophysics-UOW.

## Acknowledgements

The authors would like to thank members of the Spenkelink, van Oijen, and Griffiths labs for helpful discussions. This work was supported by the Australian National Health and Medical Research Council (Investigator grant 2007778 to L.M.S.), the Human Frontiers of Science Program (Program grant RGP0021/2020 to A.D.G. and A.M.v.O.), the Australian Research Council (Discovery Project Grant DP240101399 to L.M.S., A.M.v.O., and A.D.G., Centre of Excellence in Quantum Biotechnology CE230100021 to H.Y.

## Contributions

**S.H.M.**: conceptualisation, methodology, software, validation, formal analysis, investigation, data curation, writing – original draft, visualisation. **D.M.G.**: formal analysis, investigation. **H.J.P.**: methodology, formal analysis, investigation, visualisation. **A.C.F.**: investigation. **H.Y.**: methodology, formal analysis, writing – original draft, supervision. **M.R.-C.**: conceptualisation, methodology. **A.D.G.**: conceptualisation, supervision, project administration, funding acquisition. **A.M.v.O.**: conceptualisation, supervision, project administration, funding acquisition. **L.M.S.**: conceptualisation, methodology, validation, formal analysis, investigation, data curation, writing – original draft, supervision, project administration, funding acquisition.

## Ethics declarations

### Competing interests

This work was financially supported by Quantum-Si. Antoine van Oijen and Andrew Griffiths were members of the Quantum-Si Scientific Advisory Board.

## Extended Data

**Extended Data Fig. 1.**
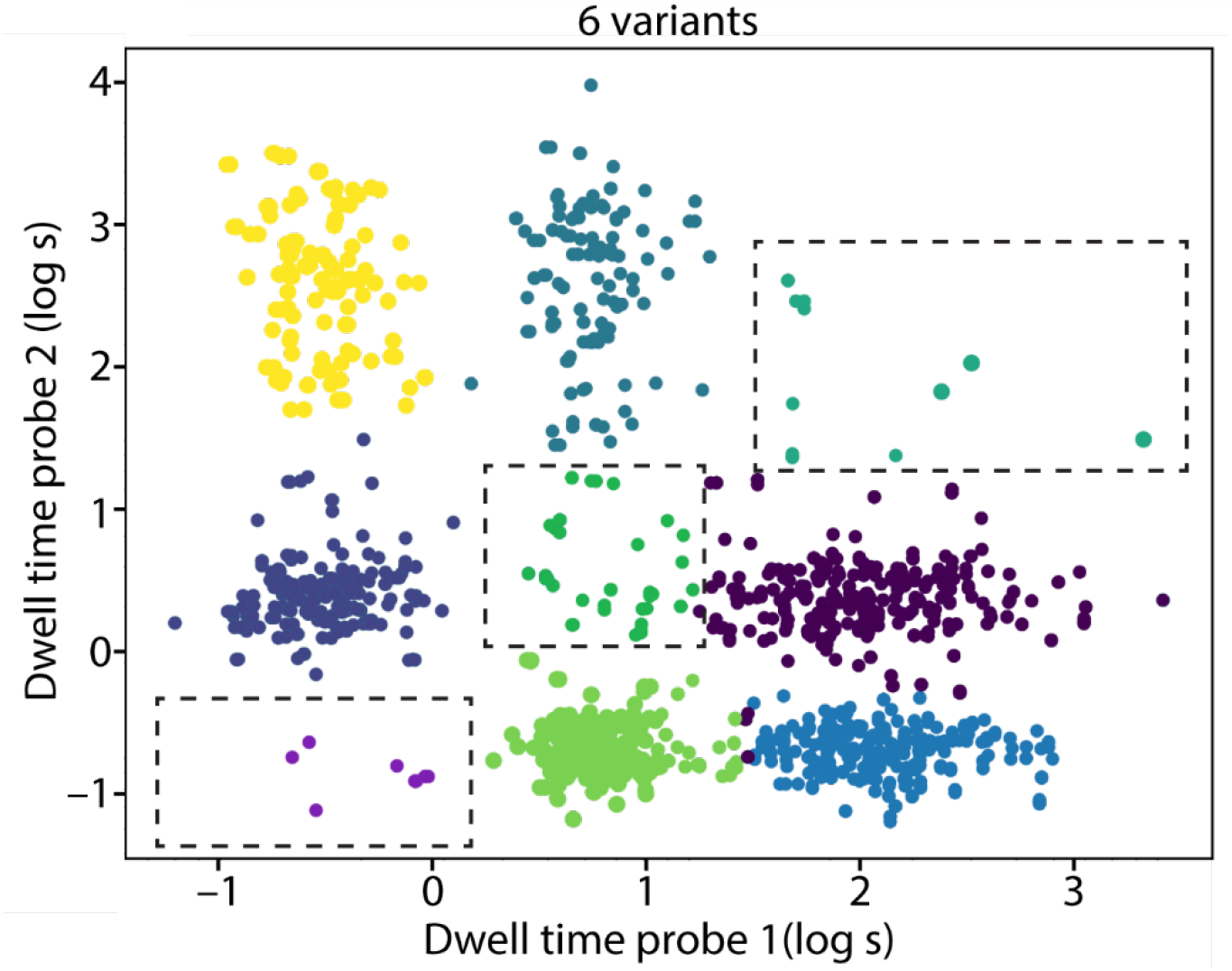
Quantification of accuracy. We measured classification accuracy using six two-index barcodes, instead of nine (Fig. 2), excluding barcodes with both indices of the same length (7nt–7nt, 8nt–8nt, and 9nt–9nt; dashed boxes). Each dot in the graph represents a single two-index barcode. Each colour represents a cluster of identical barcodes. Clusters are defined to be 1.5 standard deviations around the cluster means (Extended Data table 2). Any dot appearing in the dashed boxes represents a false call. Based on this, we observed an accuracy exceeding 95% (*e.g*. 5% falls calls).

**Extended Data Fig. 2.**
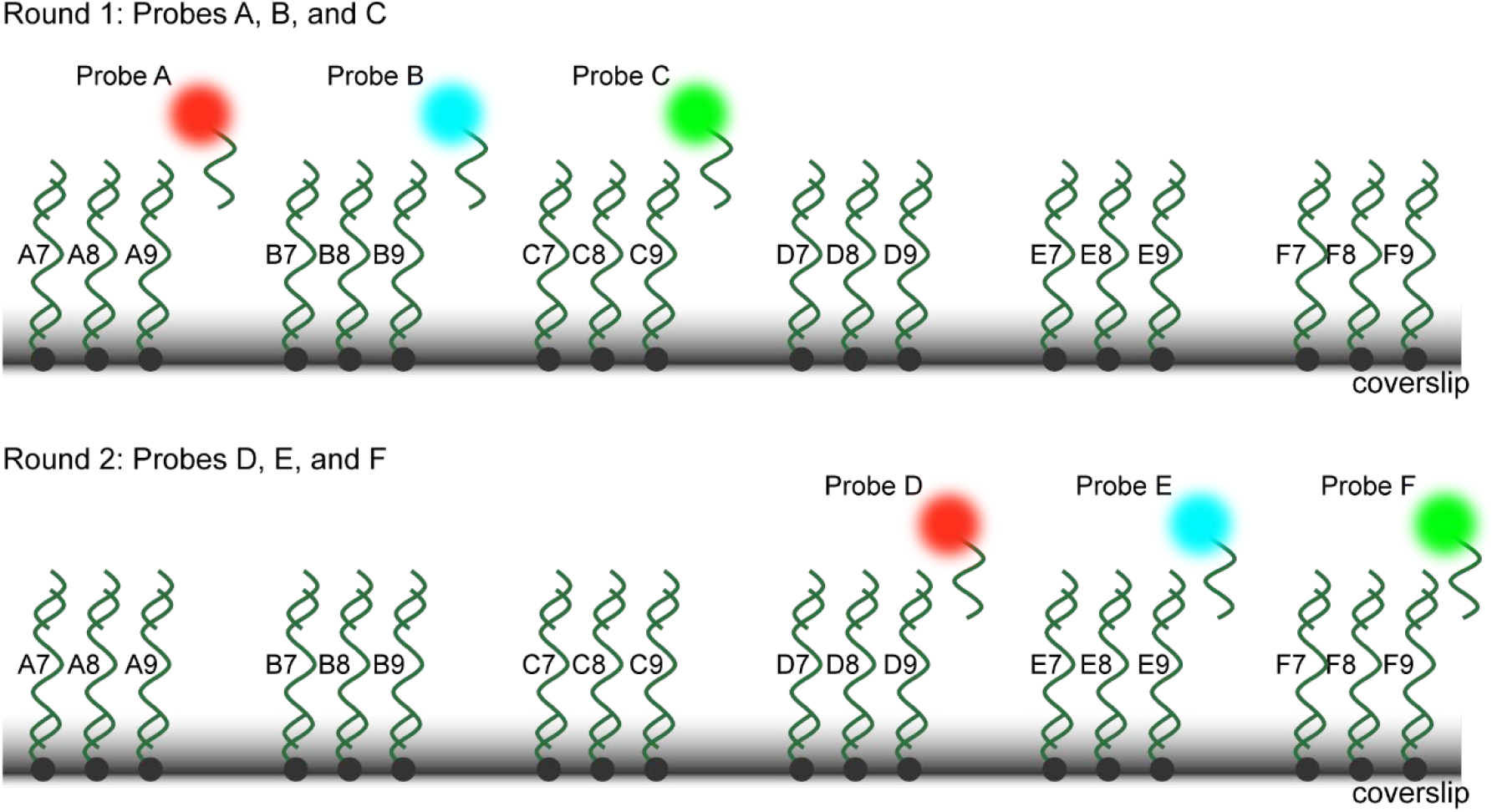
Measurement of 18 unique barcodes using six orthogonal probes. We generated a set of 18 indices, by combining six orthogonal probes (A–F), each capable of binding three distinct single-index DNA barcodes with 7-nt, 8-nt, or 9-nt complementarity to the probe. All 18 barcodes are immobilised on the microscope coverslip simultaneously. Probes A–C are imaged in the first round (top panel), and probes D–F in the second round (bottom panel). By using three spectrally distinct fluorophores (two probes per colour) and resolving binding kinetics for each probe, we generate 6 × 3 = 18 unique indices.

**Extended Data Fig. 3.**
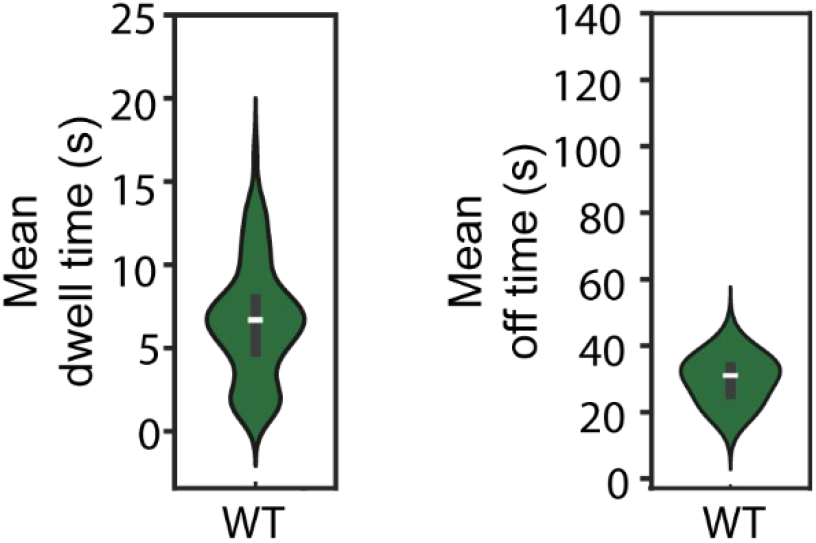
Measurement of wild-type ClpS in isolation. A measurement of the peptide binding kinetics in an experiment that contained only wild-type ClpS, reproduce the distributions of mean dwell times (left) and mean off times obtained in Fig. 3c.

**Extended Data Fig. 4.**
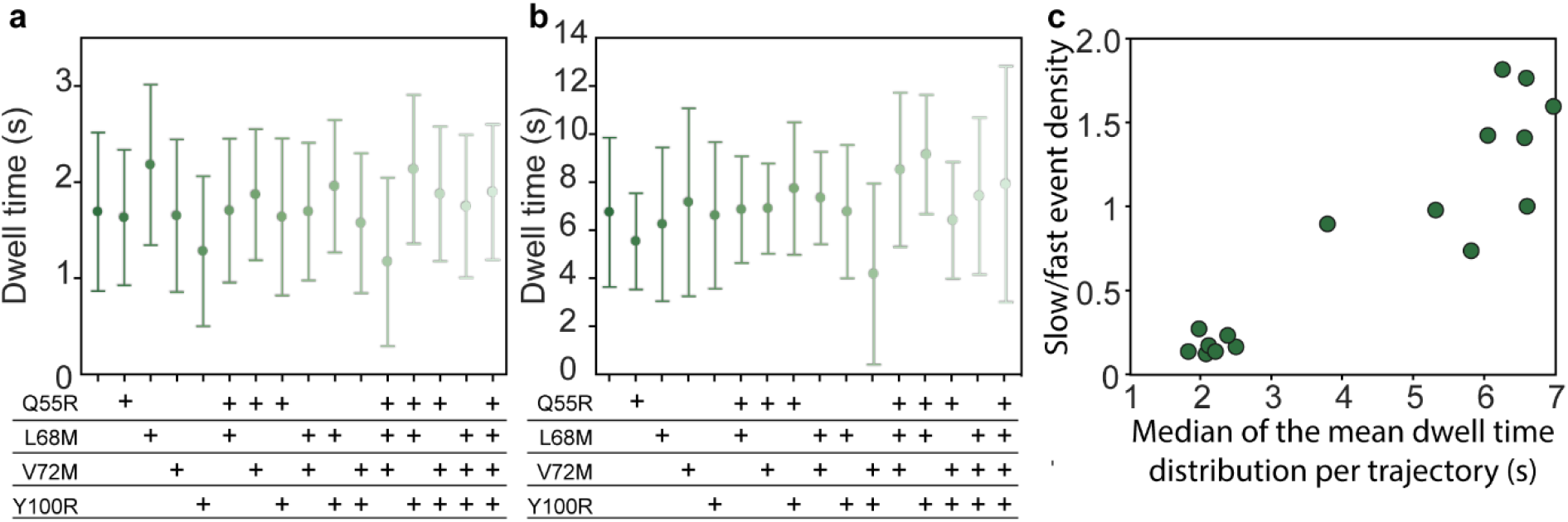
Quantification of the bimodal dwell distributions. The violin plots for the dwell time distributions shown in Fig. 3e were fit with the sum of two Gaussians to calculate **a**, the mean dwell time of the fast population for each variant (∼1.5 s for all variants), and **b**, the mean dwell time of the slow population for each variant (∼7s for all variants). Error bars represent the standard deviation. **c**, the ratio of the densities for the slow and fast populations *vs* the median of the mean dwell time distributions per trajectory, shows that the dwell time is largely governed by the ratio of the two kinetic states.

**Extended Data Fig. 5.**
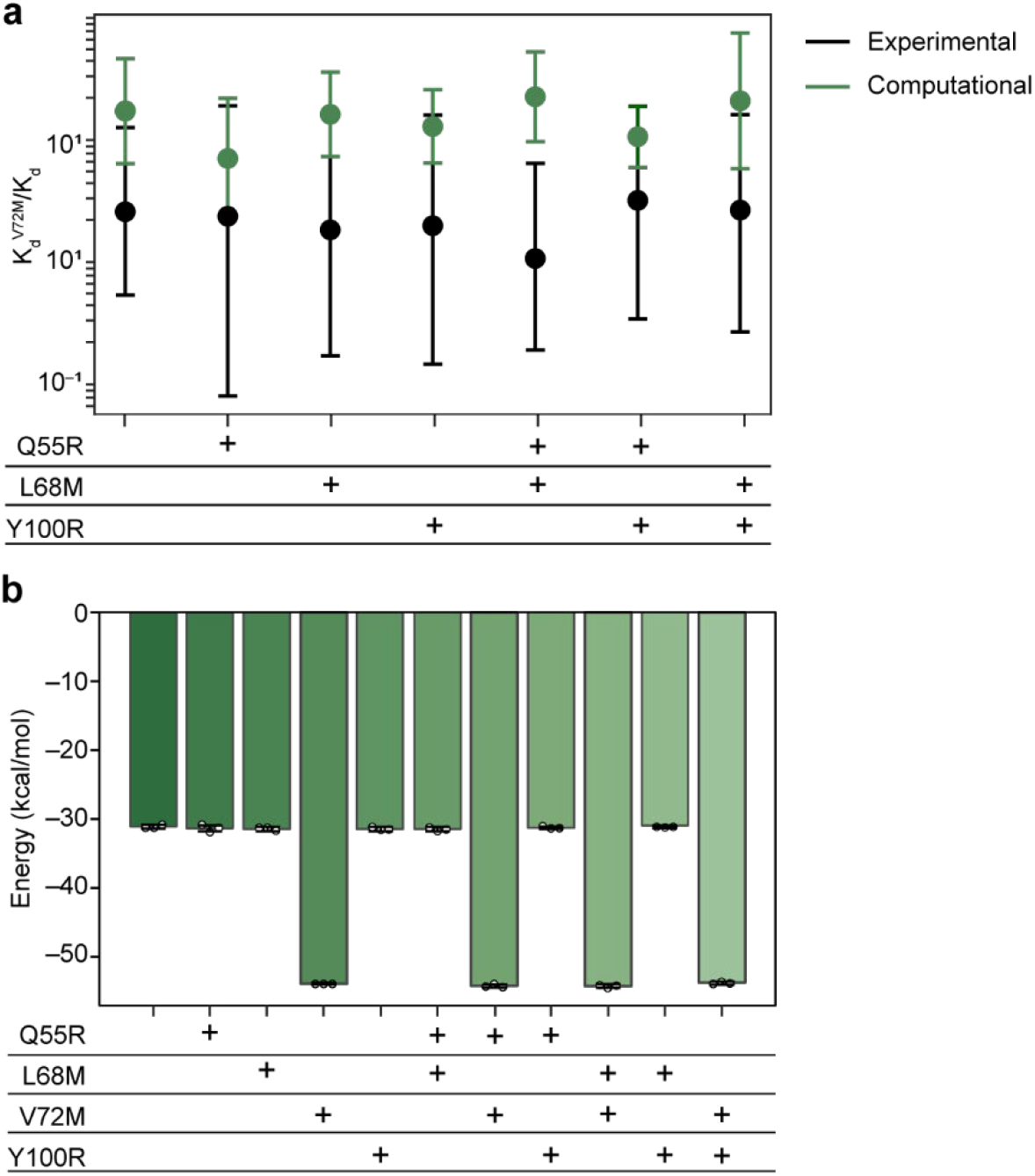
Molecular dynamics simulations show effect of V72M. **a**, Ratio of *K*_d_ for V72M-Containing to those for Non-V72M-Containing Mutations. Assessing the Effect of the V72M Mutation on the Dissociation Constant. The experimental data is shown in black, with the error bars representing the range of the K_d_^+V72M^/K_d_ quantity. The simulated K_d_^+V72M^/K_d_ is shown in green, and the error bars represent the standard deviation of the TI replicates. **b**, Electrostatic interaction energies of the terminal leucine residue with ClpS Val/Met72. This figure depicts the component of the interaction energy between the ClpS residue 72 (which can be either Val or Met depending on the variant) and the N-terminal leucine residue of the peptide. Each data point (the circles on the bars) is the average of this component of the interaction energy across the 50,000 trajectory frames of a single replicate calculated with the enedecomp module of CPPTRAJ. The bars represent the means of the triplicate data, and the error bars represent the standard deviations. There is a significant decrease in energy with mutation from the wild-type valine to methionine. This decrease in energy is not affected by additional mutations.

**Extended Data Fig. 6.**
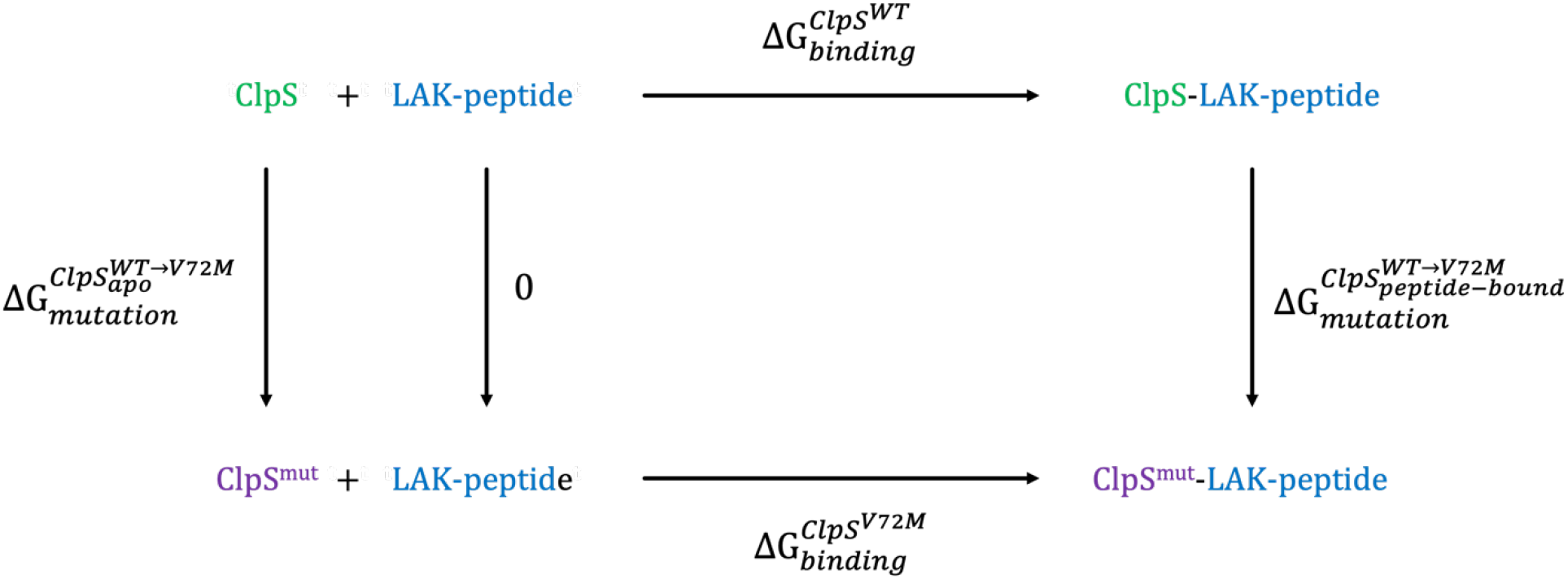
Thermodynamic cycle used in alchemical thermodynamic integration simulations. Wild-type ClpS is transformed to the variant ClpS^V72M^ in apo and in the peptide-bound state. The simulations involve the vertical alchemical transformations.

**Extended Data Table 1.**
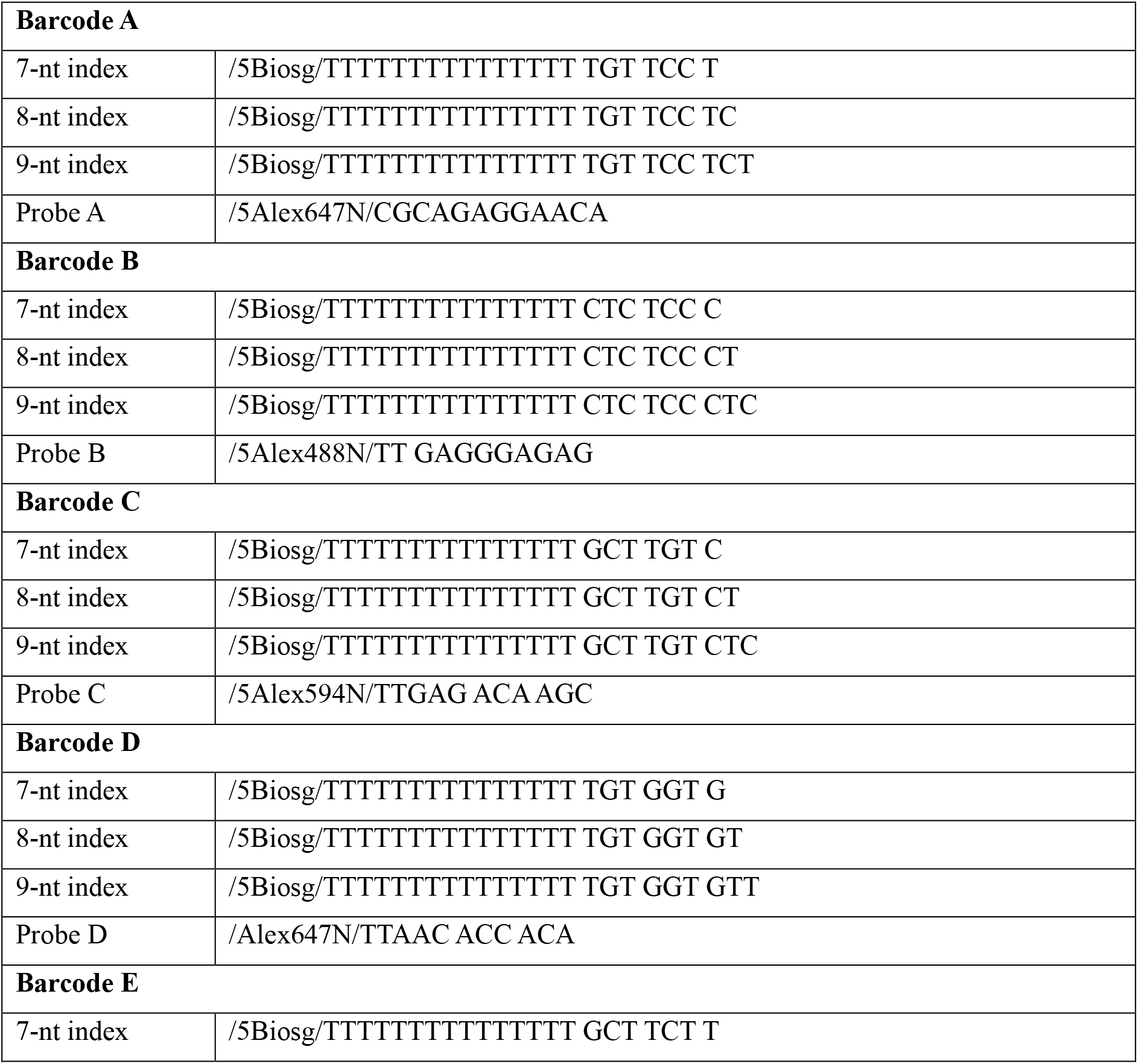
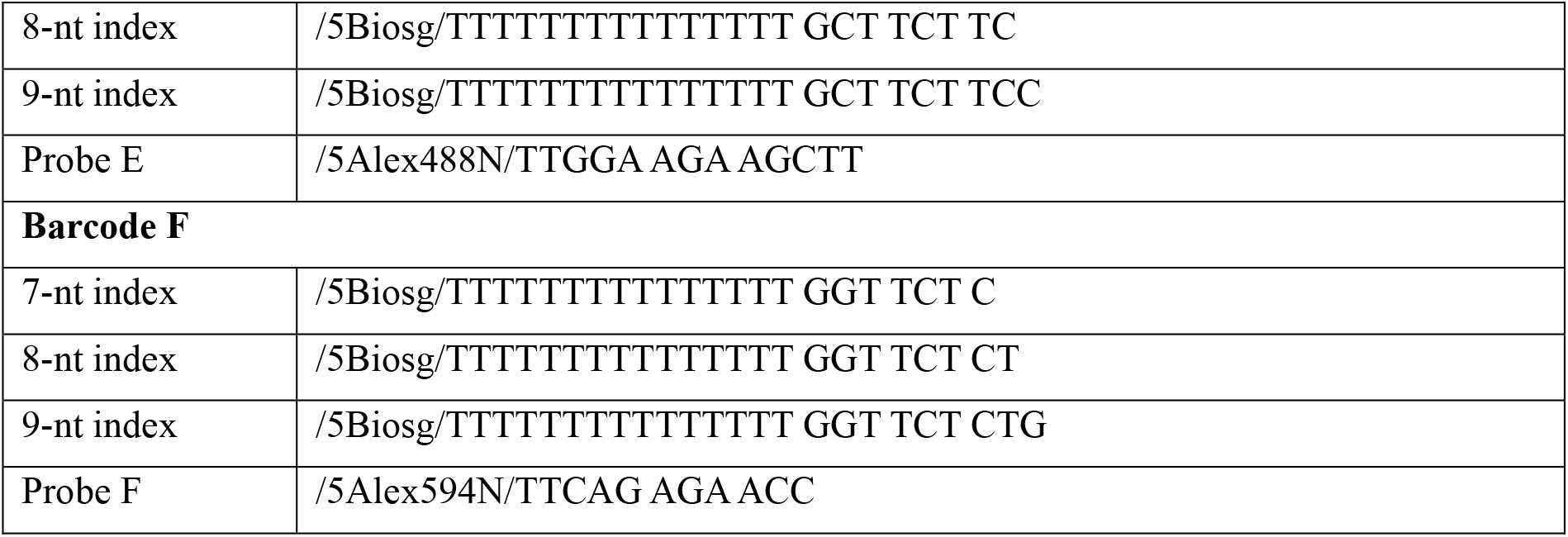
Index and probe sequences

**Extended Data Table 2.** Dwell times (s) for all indices.

|  |  | Index length (nt) |  |  |
| --- | --- | --- | --- | --- |
| Barcode |  | 7 | 8 | 9 |
|  | A | 0.34 ± 0.05 | 2.39 ± 0.02 | 8.01 ± 0.44 |
|  | B | 0.46 ± 0.05 | 1.44 ± 0.08 | 20.43 ± 1.39 |
|  | C | 1.40 ± 0.30 | 7.70 ± 2.20 | 38.10 ± 5.90 |
|  | D | 0.58 ± 0.03 | 3.01 ± 0.13 | 20.37 ± 0.28 |
|  | E | 2.70 ± 0.20 | 9.70 ± 0.08 | 74.73 ± 17.08 |
|  | F | 3.00 ± 1.50 | 9.00 ± 2.70 | 23.89 ± 2.29 |

**Extended Data Table 3:**
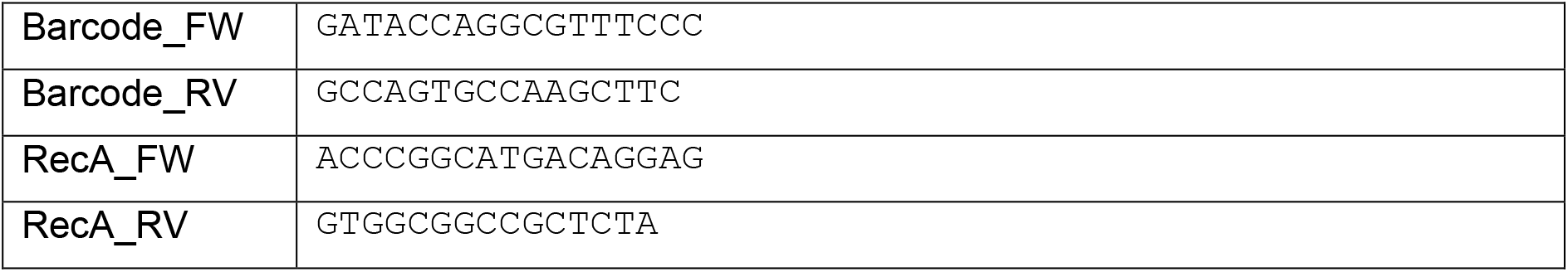
PCR primers for SNAP–ClpS constructs

**Extended Data Table 4:**
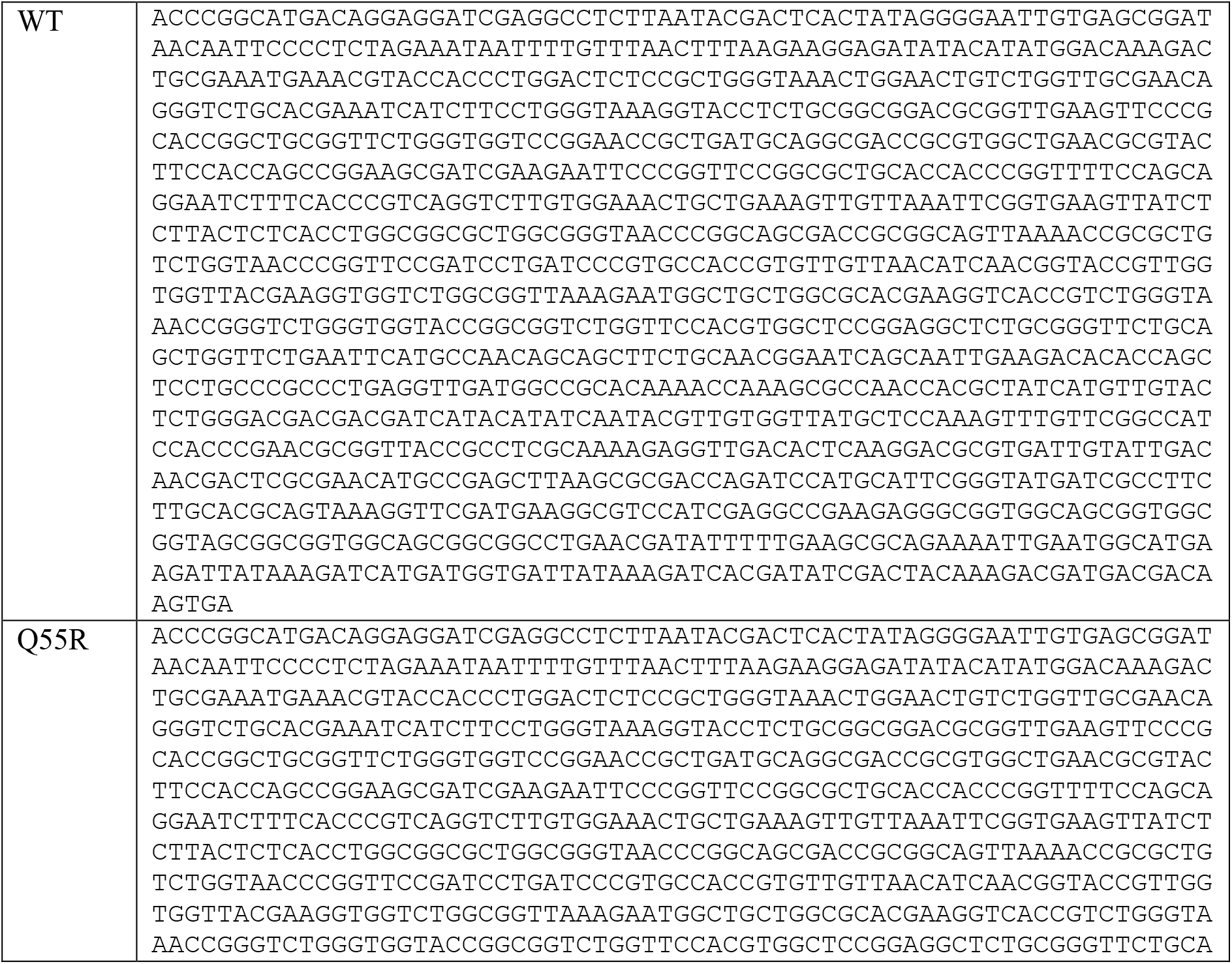
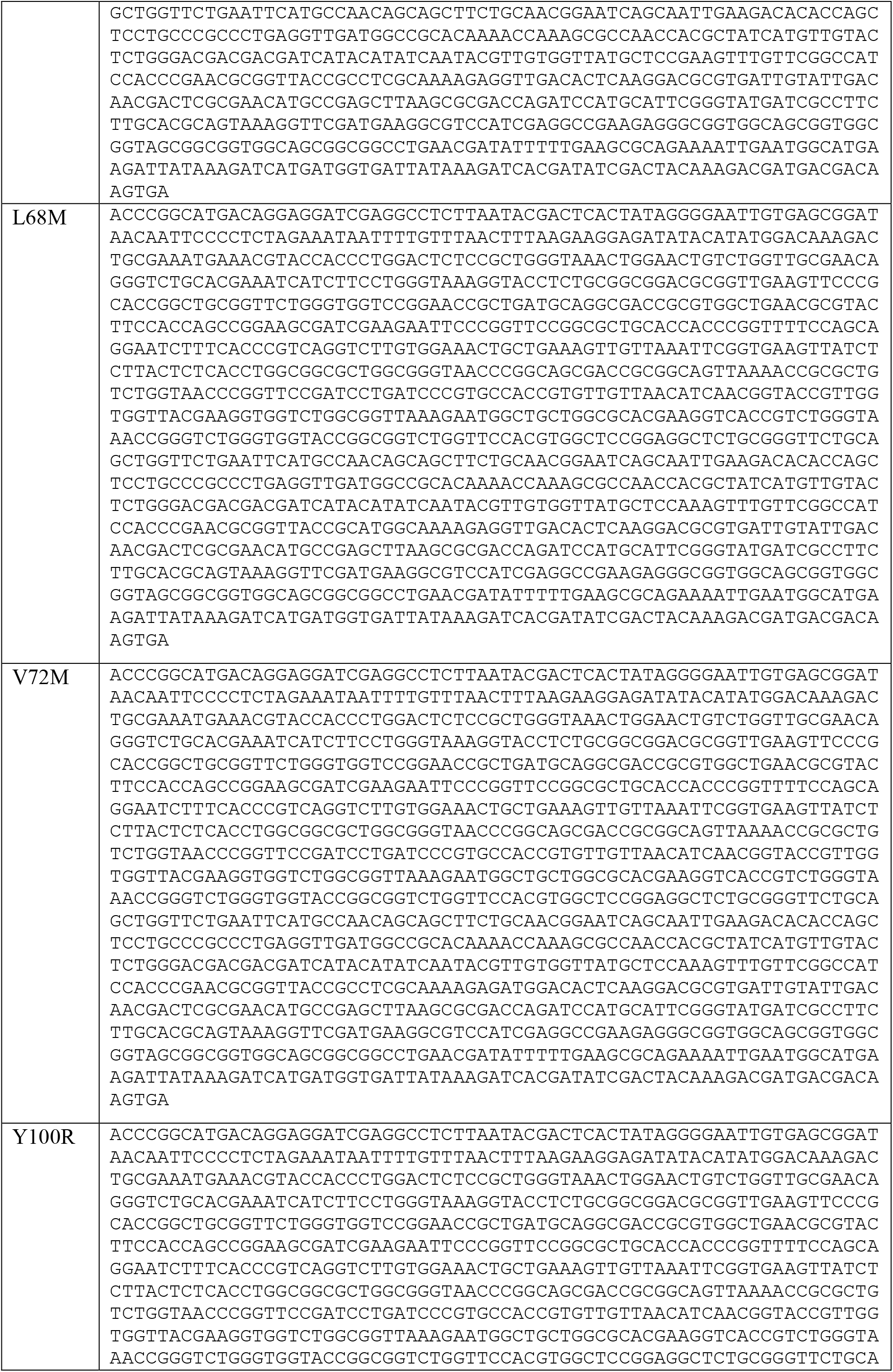
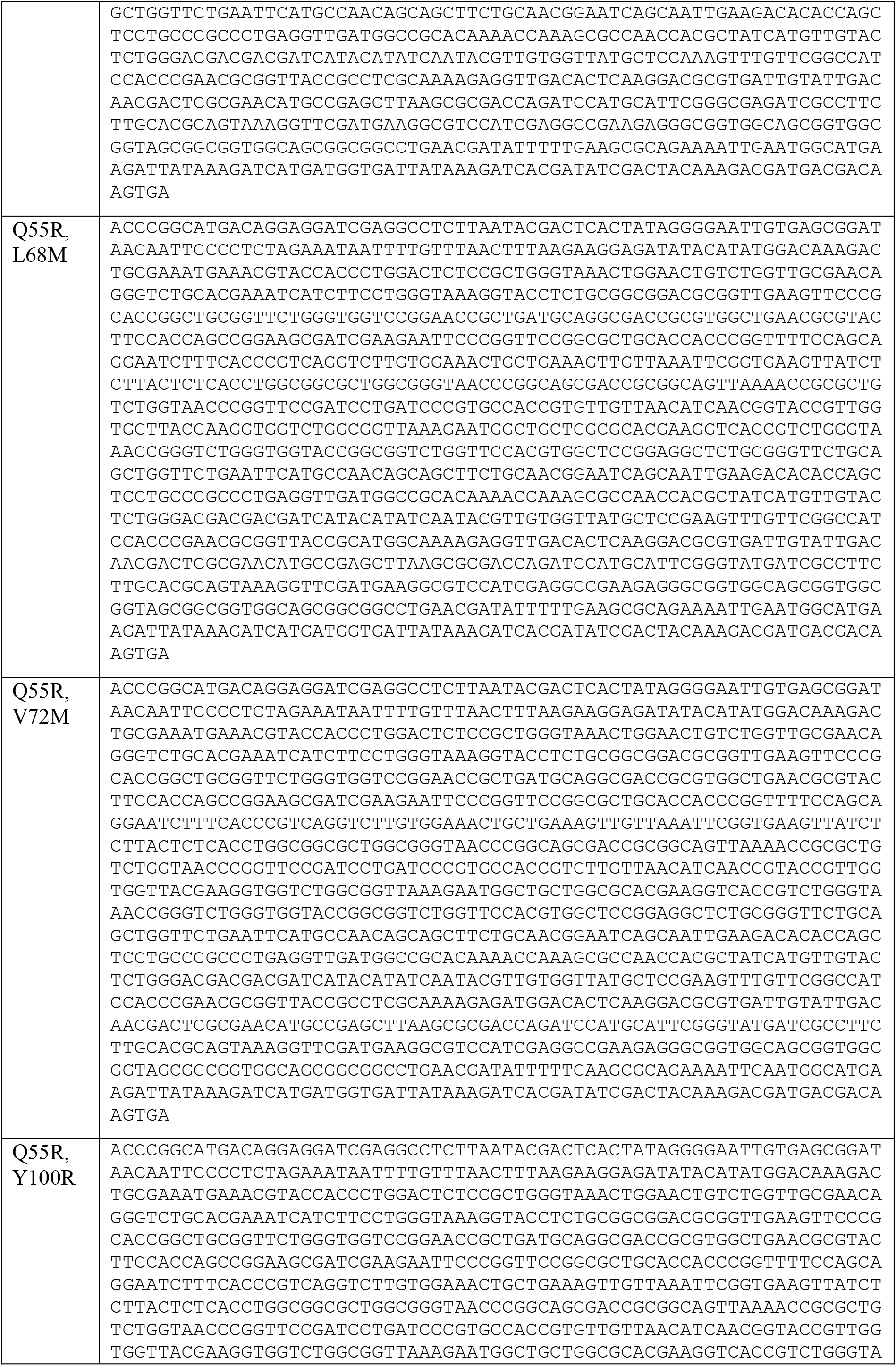
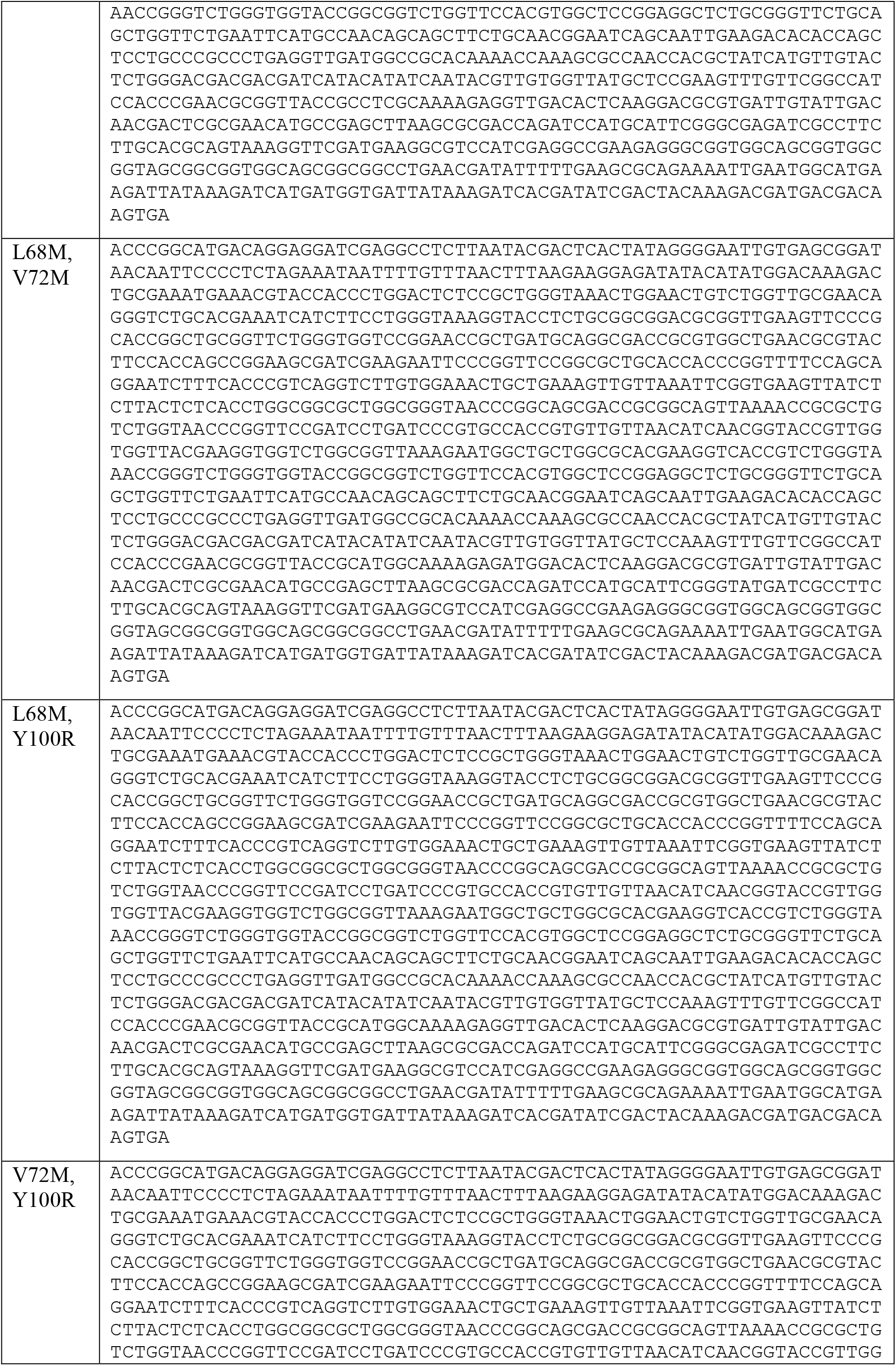
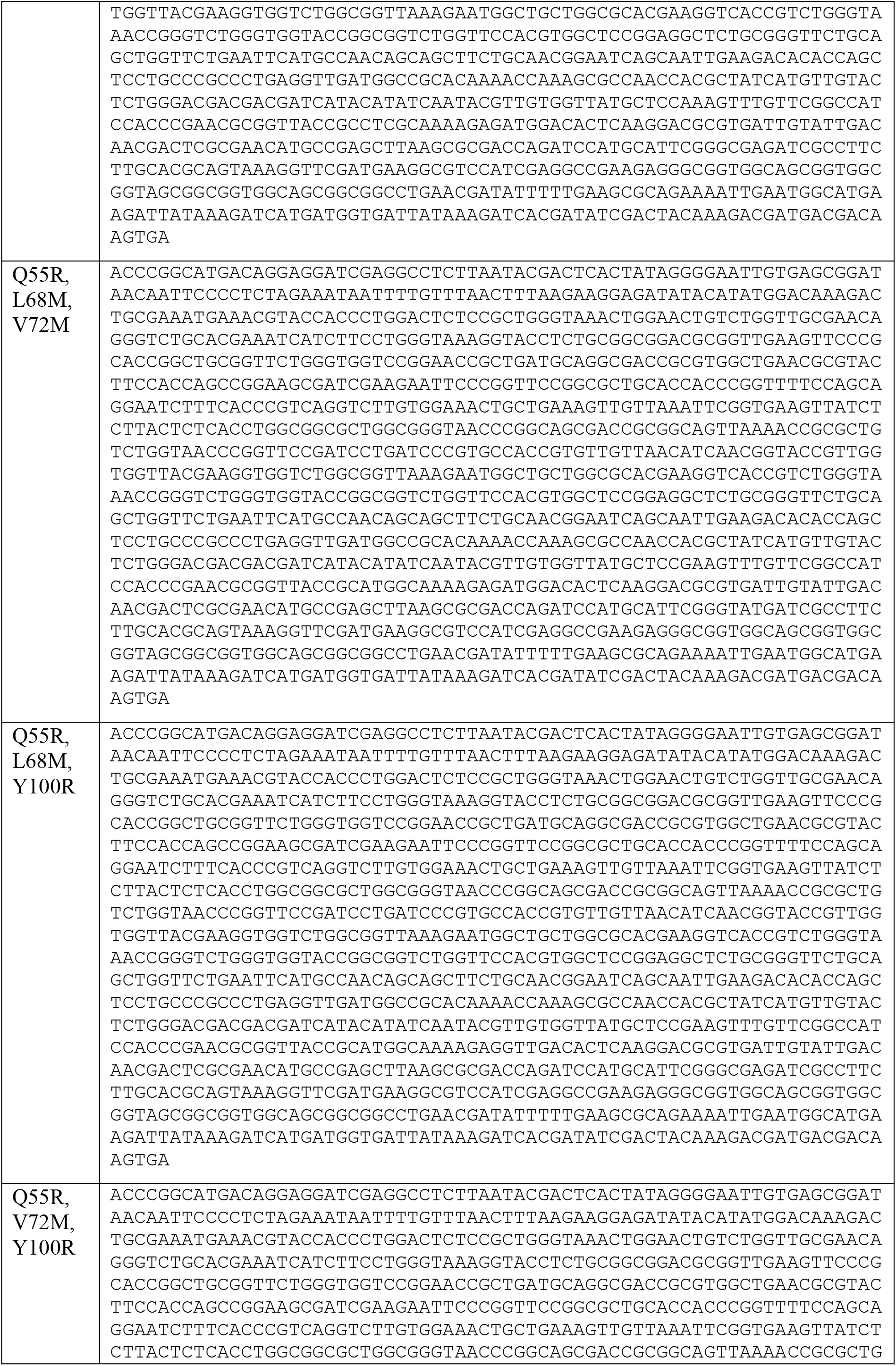
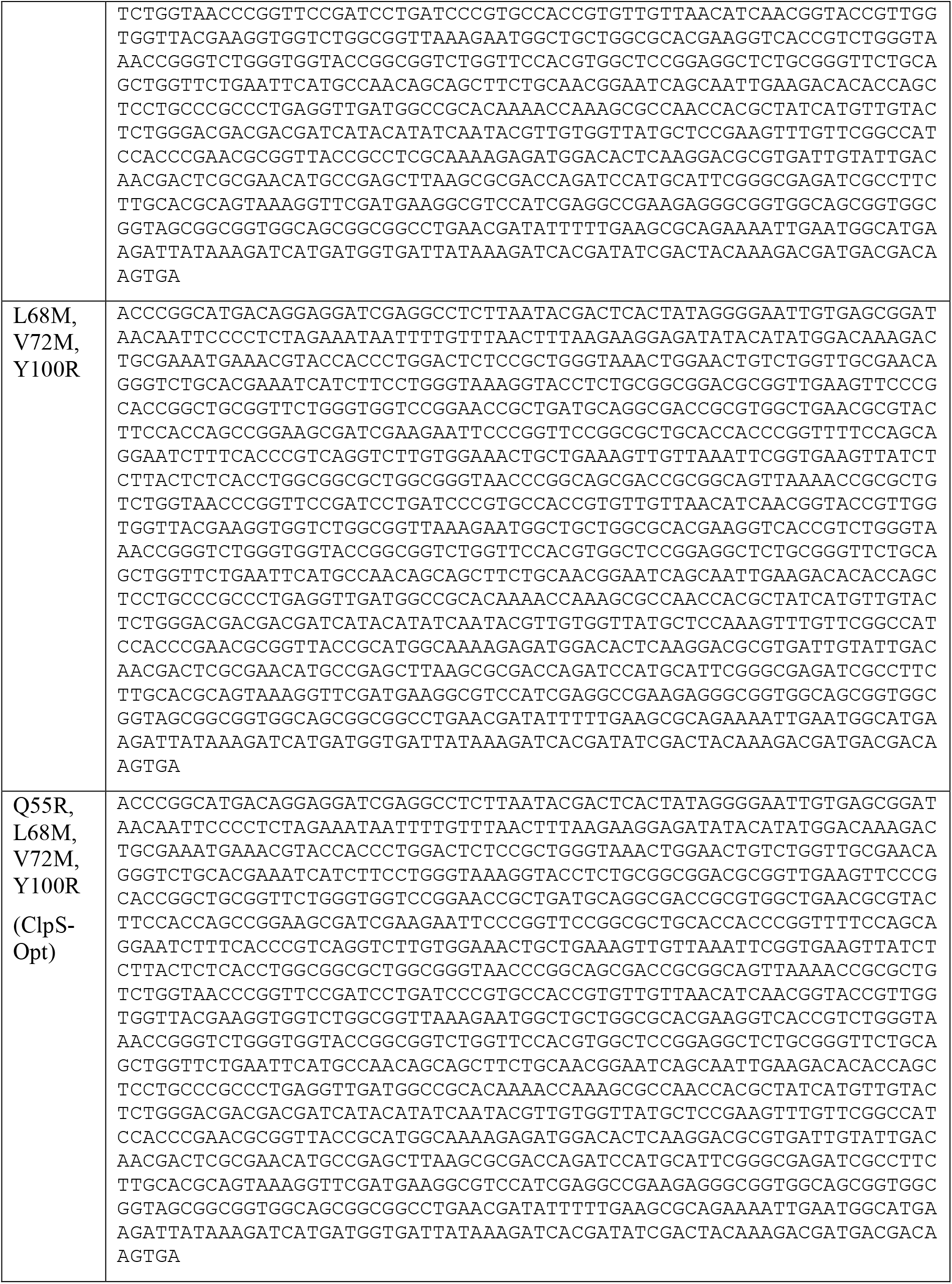
SNAP–ClpS constructs

